# Regeneration of Starfish Radial Nerve Cord restores animal mobility and unveils a new coelomocyte population

**DOI:** 10.1101/2022.10.24.513254

**Authors:** Filipe Magalhães, Claúdia Andrade, Beatriz Simões, Fredi Brigham, Ruben Valente, Pedro Martinez, José Rino, Michela Sugni, Ana Varela Coelho

## Abstract

The potential to regenerate a damaged body part is expressed to a different extent in animals. Echinoderms, in particular starfish, are known for their outstanding potential to regenerate cell, tissue, organ, and body parts. For instance, humans have restricted abilities to restore organ systems being dependent on limited sources of stem cells. In particular, the potential to regenerate the central nervous system is extremely limited, explaining the lack of natural mechanisms that could overcome the development of neurodegenerative diseases and the presence of traumatisms. Therefore, understanding the molecular and cellular mechanisms of regeneration in starfish could lead to the development of new therapeutic approaches in humans. In this study, we tackle the problem of starfish central nervous system regeneration by examining anatomical and behavioral traits, including external anatomic anomalies, the dynamics of coelomocytes populations and neuronal tissue architecture. We noticed that several anatomic anomalies were evident and detected that the injured arm could not be used anymore to lead the starfish movement. Those seem to be related to defense mechanisms and protection of the wound. In particular, histology showed that tissue patterns during regeneration resemble those described in holothurians and in starfish arm tip regeneration. Flow cytometry coupled with imaging flow cytometry unveiled a new coelomocyte population during the late phase of the regeneration process. Morphotypes of previously characterized coelomocytes populations were described based on IFC data. Further studies of this new coelomocyte population might provide insights on their involvement in radial nerve cord regeneration.

## Introduction

Regeneration is an intrinsically conservative post-embryonic developmental process that repairs and replaces cells, tissues, organs, and body parts of a given organism, and it is expressed to a different extent among all animal phyla (Alvarado and Tsonis 2006; Candia Carnevali 2006). New cells develop in the established context of mature injured tissues depending on their histogenetic and morphogenetic plasticity (Ben Khadra et al. 2017). Regeneration is crucial for echinoderms survival and provides a necessary programed complement for asexual reproduction (Byrne 2020; Candia Carnevali and Burighel 2010). Particularly, members of the Asteroidea, Ophiuroidea and Holothuroidea are all well known for their outstanding potential to regenerate the CNS, as well other tissues or body parts (Allievi et al. 2022; Ben Khadra et al. 2015a; Ben Khadra et al. 2015b; Ben Khadra et al. 2017; Ben Khadra et al. 2018; Franco et al. 2012; Franco et al. 2014). Critically, neural structures are the first to be regenerated in stellate echinoderms, a feature that underlines their fundamental regulatory role in these animals’ regeneration (Ben Khadra et al, 2018). In adult starfish, the cellular mechanisms that underlie the regeneration of a new tissue are dedifferentiation, transdifferentiation, and/or migration of cells derived from local tissues (Alvarado and Tsonis, 2006; Mashanov et al. 2017; Ferrario et al. 2020). Up to date studies specifically on the regeneration of the CNS have been mainly performed in holothuroids. According to the literature, in these animal models, after CNS injury radial glial cells are activated, undergo dedifferentiation and give rise to new glial and neuron cells, reconstituting completely the nerve morphology (Mashanov et al. 2013). Although already present in injured sea cucumber CNS, proliferation of these radial glial cells increases during neural regeneration (Mashanov and Zueva 2019). In addition to glial cells, the contribution of migratory cells from more distant regions was also described, although their nature remains to be clarified. It has been hypothesized that this latter process mainly relies on the reprogramming of adult differentiated cells instead of the recruitment of adult undifferentiated cells (Mashanov 2017). Remarkably, Zheng et al. (Zheng et al. 2022) highlighted that starfish larvae (*Patiria miniata*) regenerate their nervous system by re-specification of existing neurons. Specifically, in the larval stage of this starfish, injured neurons expressed the gene sox2, which leads these cells to re-enter neurogenesis and form new differentiated neurons. This regulatory gene is also involved in human neuronal regeneration, where it has a critical role in the maintenance of both embryonic and neural stem cells, where it has a critical role in the maintenance of both embryonic and neural stem cells (Pagin et al. 2021).

The neuro-regeneration in starfish displays some common features to mammals, such as the regulation of nerve cord protein phosphorylation (Franco et al. 2012), the presence of pluripotency factors orthologs with Yamanaka factors of the mammalian cells (Mashanov et al. 2015), the expression of Hox gene homologs (Khadrha et al. 2014), the action of growth factors, neuropeptides, and neurotransmitters (Thorndyke and Candia Carnevali 2001), and of related proteins with relevant roles (Franco et al. 2014). Additionally, to the neuro-regeneration similarities, the close taxonomic proximity and the similar features, development and neural communication of their CNS make from Asteroidea insightful animal models for the study of CNS regeneration mechanisms, namely envisaging the development of new therapeutic approaches.

Among other cell types known to be involved in the regenerative processes, the coelomocytes also play a key role in starfish models, since they are among the first cells to be mobilized to the site of injury protecting the inner environment from foreign material, healing the wound, and initiating the restoration of the missing structures (Pinsino et al. 2007; Smith et al. 2018; Khadra et al. 2017). Different populations regulate different process during these first reaction phases. Thrombocytes-like cells have a similar role to the human platelets when the starfish suffers an injury. These cells are involved in hemostasis, migrating to the site of the lesion and forming a clot that separates the inner from the outer environment to prevent bleeding (Ben Khadra et al. 2015). The phagocytes, besides phagocytic activity against pathogens, form a network at the site of the lesion supporting the clot formation by the thrombocytes-like cells (Andrade et al. 2021; Khadra et al. 2017). Coelomocytes could also be involved in wound repair since they show an increasing level of Hsp70 during this process, and this molecule induces the release of pro-inflammatory cytokines during repair phase in mammals (Ben Khadra et al. 2017; Pinsino et al. 2007).

With the purpose to give new insights into starfish neuronal regeneration, we implemented, as herein described, several experiments focusing only on the study of the CNS regeneration triggered by a traumatic injury of the radial nerve cord (RNC) in *Marthasterias glacialis* (Linnaeus, 1758). Behavioral assays and external anatomical observations were performed to acknowledge the constraints of the nerve lesion in the locomotion and external morphology of starfish. Histological analyses were used to study the cell-tissue pattern during the regeneration of the injured nerve tract. In addition, the role of coelomocytes was assessed by flow cytometry (FC) coupled with imaging flow cytometry (IFC) following the implemented experimental protocol (Andrade et al. 2021). The objective being to characterize the dynamics of the coelomocyte populations circulating in the coelomic fluid during the regenerative process. Our results give a new perspective of the cellular mechanisms behind nerve regeneration and suggests new roles for a new coelomocyte population involved in this process.

## M&M

### Animal collection and maintenance

Adult specimens of the starfish *M. glacialis* (diameter between 14 and 32 cm), without previous signs of regeneration and considered healthy, were collected at low tide on the west coast of Portugal (Estoril, Cascais), outside the breeding season. The animals were placed in captivity at the “Vasco da Gama ‘‘ Aquarium (Dafundo, Oeiras), in open-circuit tanks with re-circulating seawater, with a temperature of 15°C and salinity of 33‰. They were fed *ad libitium* on a mussel diet. All specimens were maintained in the same conditions throughout the whole experimental period to avoid variability due to abiotic factors.

### Induction of regeneration

Prior to induction of regeneration, starfish were anesthetized by immersion in 4% (w/v) magnesium chloride in seawater. After the individuals were relaxed, induction of nerve regeneration was triggered by the partial excision of around 1 cm of the radial nerve close to the tip of the arm (at a position around 1 cm from the arm tip). The RNC is “external” and easily accessible from the oral side. The arm immediately to the right of the madreporite was considered as arm 1 and the following ones were counted in the anticlockwise direction. For each animal, the nerve was partially excised in two adjacent arms (1 and 5) for the behavioral assay and only in arm 1 for cytometry and histology experiments. Consequently, two classes of arms were considered for each animal: the arms to which a partial excision of the nerve was done, i.e., those with the regenerating nerve (RN) and those with the intact nerve (NRN).

### Animal displacement and external anatomy observation

Before and after nerve ablation, an animal displacement assay was performed by placing the starfish in the center of an empty tank with seawater. For 2 minutes, the number of times each arm was used to direct the animal movement in the tank (i.e., as the leading arm of the movement) was registered. The percentage of times that each arm was used as a leading arm was obtained by dividing this value by the total number of movements. This assay was performed before nerve ablation, and at days 1, 4, 8, and 14 post-nerve ablation (PA). A t-test was carried out to compare the level of use between the two arm classes for each time-point. Furthermore, at 1 hour, and days 1, 2, 4 and 8 PA, visual inspection of the external anatomy was made at the site of injury, and several parameters were monitored: a) ambulacral feet movement or retraction, b) presence of external edema, c) irregularities in the spicules orientation along each arm and d) bending of the arm at the injury site.

### Coelomic Fluid collection

The study of coelomocytes populations and nerve morphology during the initial phase of the RNC regeneration process was carried out over three-time points (1 day, 7 days, and 14 days PA). At each time point, coelomic fluid was collected for FC and IFC analysis from the two arms where nerve regeneration was induced. Coelomic fluid was harvested through perforation of the epidermis at the tip of the starfish arm, to avoid the disruption of internal organs, with a 21-gauge-butterfly needle. By gravity, the fluid was directly transferred to a Falcon tube. Subsequently the tip (approximately the last third of the whole arm length) of these same arms, including the nerve amputation zone, were cut with a scalpel for histological analyses. Tissues were fixed in Bouin solution, renewed each week, and stored at 4° C, until paraffin embedding.

### Histology of starfish arm tissues

Bouin fixed samples were processed by standard histological protocol, as described in Ben Khadra et al. (2015b). Briefly, after several washes in tap water the samples were dehydrated in an increasing ethanol series, cleared in xylene, washed in a 1:1 xylene and paraffin solution and then embedded in pure melted paraffin (56-58°C). Sagittal/parasagittal sections (7-10 μm thickness) were prepared with a Leitz 1512 rotary microtome. Two or 3 sample slices were placed on top of a glass slide. Those serving the purpose of this study (i.e., including the nerve gap or the total nerve tissue) were stained following Milligan’s Trichrome procedure (Milligan 1946). Stained sections were mounted with Eukitt® (5% acrylic resin and 55% xylene mounting medium), observed and photographed under a Jenaval light microscope provided with a Leica EC3 Camera and processed using the Leica Application Suite LAS EZ Software (Version 1.8.0).

### Flow cytometry (FC)

CF was resuspended with a micropipette and was preliminary filtered through a 40 μm mesh to remove cellular aggregates and debris. To discriminate viable coelomocytes from cellular debris, 0,25μL of 5 μg/mL DRAQ5 (Invitrogen) was added to 1 mL of each sample. DRAQ5 is a membrane permeable DNA dye that allows the detection of live cells (Smith et al., 2004). Samples were run through BD FACSAria™ III (BD Biosciences) flow cytometer, using a 620/20 BP filter and a laser of 633 nm. The data obtained by the FC was subsequently analyzed using the software FlowJo (version 10.8.1, Becton, Dickinson & Company). The gating strategy is described in **Supplementary Fig.1**.

### Imaging flow cytometry (IFC)

Samples were treated using the protocol described for FC analysis. Coelomocyte images were acquired using the INSPIRE software of the ImageStreamX Mark II imaging flow cytometer (Luminex Corporation, Austin, TX) at the “Instituto de Medicina Molecular João Lobo Antunes, Faculdade de Medicina”, University of Lisbon, Portugal. Cells were imaged at 60x magnification using a 642 nm laser beam at 150 mW for DRAQ5 excitation with a detection window of 642-745 nm (channel 6). Brightfield images were acquired in channels 1 and 9. Data were analyzed, and a characterization of each coelomocyte subpopulation was made based on specific morphologic characteristics resorting to the software IDEAS (version 6.2, Luminex Corporation, Austin, TX).

## Results

### Animal displacement recovery and external anatomic anomalies induced by nerve ablation

To evaluate the recovery of mobility after injury, the percentage of times that a RN or a NRN arm was used as the leading one during starfish movement at each time-point, was determined. The t-test results in the comparison of use between RN and NRN arm-classes over time are depicted in **Table 1 (**and supplementary figure 1). Before induction of regeneration, all RN or NRN arms were indistinctly used by the starfish as leading arms. After nerve ablation significant differences (p<0.05) were found in the use of leading arms between RN and NRN classes. Our data shows that RNs are the less used arms, particularly at days 1 PA and 4 PA. At 9 PA signs RN recovery are already observed with an increment of the percentage of use that approaches that measured at time-point 0.

| Time-point<br>(day) | RN |  | NRN |  | N | p-value<br>RN vs NRN |
| --- | --- | --- | --- | --- | --- | --- |
|  | (%) |  | (%) |  |  |  |
|  | mean | s.d. | mean | s.d. |  |  |
| 0 | 43 | 25 | 57 | 25 | 14 | 0.93 |
| 1 | 15** | 19 | 85** | 18 | 15 | 0.00§ |
| 4 | 12** | 17 | 88** | 17 | 13 | 0.00§ |
| 9 | 36* | 26 | 64* | 26 | 14 | 0.00§ |
| 14 | 34* | 24 | 66* | 24 | 12 | 0.00§ |

Significant differences (\*p<0.05 or \*\*p<0.01) for the comparison between each time-point (1, 4, 9, 14 days) and time-point 0 for the total starfish movements (RN + NRN) observed, and within each arm class. § p<0.01. N-number of animals tested.

After partial excision of the RNC it was observed that in several animals some external anatomic anomalies at the injury site were evident: a) external edema, b) bending of the arm, c) irregular orientation of the spicules along each arm, and d) ambulacral feet retraction. The presence of these different features was monitored over time. An arm edema was barely visible at 1 hour PA, but it is already visible in 50% of the RNs at 1 day PA, decreasing its presence in the following days until day 8 PA. The local bending of the arm it is also noticeable 1h after nerve excision. At day 1 PA this feature is observed in the two RNs of all animals, and then it is visible for the majority of them till day 8 PA. The misalignment of the spicules showed a similar temporal pattern, appearing at 1 hour PA, being prominent at days 1 and 2 PA, and starting recovery phase at day 4 PA. Finally, the tube feet were still active at 1 hour PA, then strongly inactive at day 1 PA, and becoming totally inactive for the rest of the period analyzed, till day 8 PA (**Table 2** and **Fig 1**). This inactivity is associated with the retraction of the podia. None of these features were observed in the NRNs of all animals.

**Table 2.** Mean percentage of starfish regenerating arms with external anatomy anomalies at several time-points. For each animal the mean percentage determined for the two RN was considered.

| **Time-point** | **Edema** | **Local Arm Bending** | **Misalignment of spicules** | **Retraction of podia** |
| --- | --- | --- | --- | --- |
| **1 hour** | 6.5 | 43.0 | 72.5 | 12.0 |
| **1 day** | 50.0 | 100 | 90.0 | 80.0 |
| **2 days** | 65.0 | 96.5 | 85.5 | 100 |
| **4 days** | 36.0 | 96.5 | 79.0 | 90.0 |
| **8 days** | 17.0 | 100 | 51.5 | 100 |

**Fig 1.**
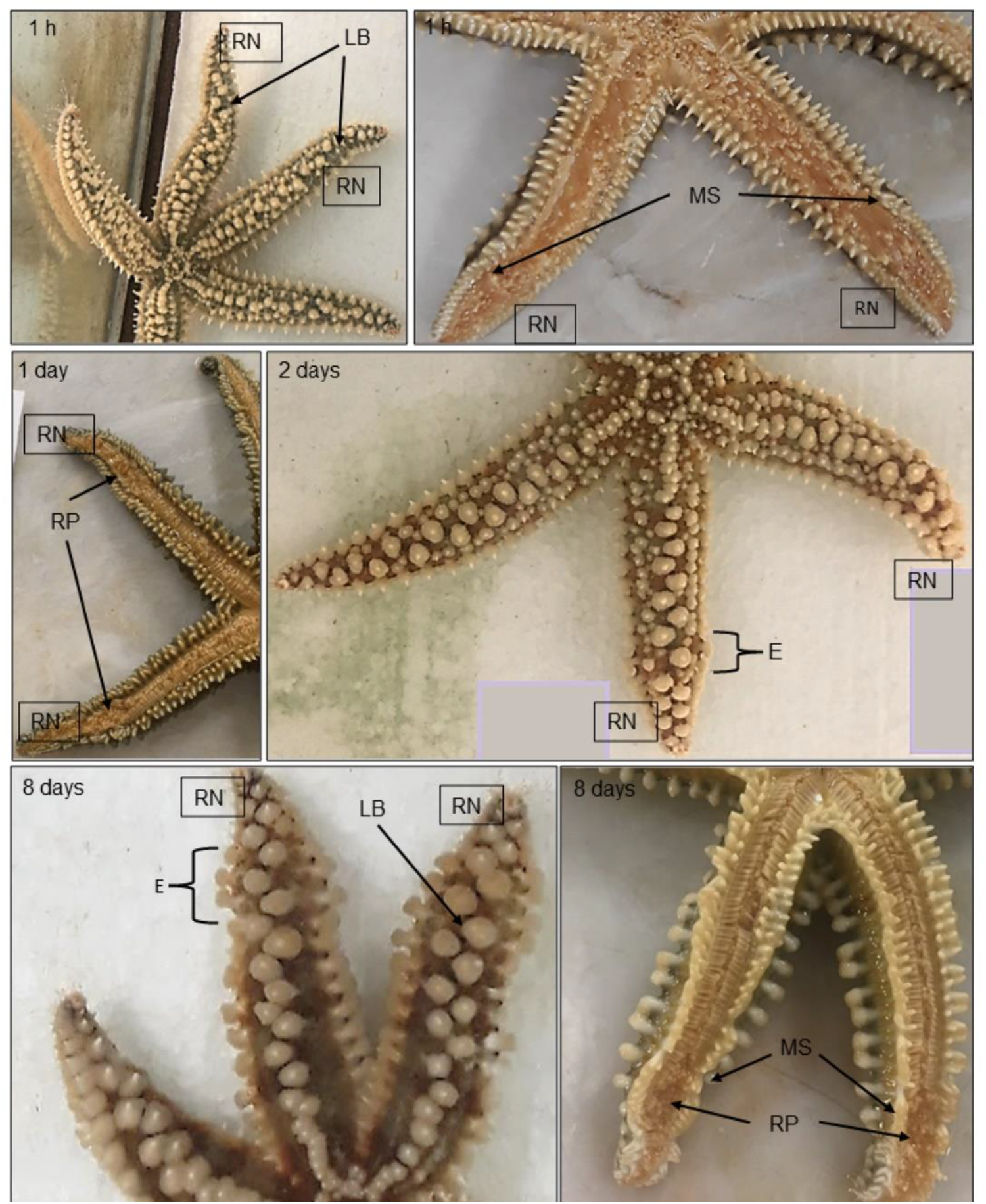
Oral and aboral view of the evolution of anatomical anomalies during RNC regeneration, at 1 hour, and days 1, 2, and 8 PA; **RN**-regenerating arm; **LB**- local bending; **MS**- misalignment of the spicules; **RP-** retracted podia; **E**- edema.

### Tissue pattern of nerve cord regeneration

The general anatomy of control uninjured nerve cords fits the description provided by Smith (1937) and it’s in line with that of other starfish species (Hymann 1955). Indeed, the RNC appears as a V-shaped structure running along the median longitudinal plane of each arm, within the ambulacral groove and laterally “protected” by two rows of podia. Differently from the other cryptosyringid echinoderms (sea urchins, brittle stars and sea cucumbers), the RNC is “external”, in direct contact with the environment (this element facilitates the surgical ablation). Above the RNC, in the aboral direction, and parallelly to it, runs the hyponeural sinus and the radial water canal, both are coelom-derived canals (**Fig 2a**). As illustrated in **Fig 2b**, the uninjured RNC is organized in two adjacent parallel bands of neuroepithelia, a thicker outer ectoneural band, and a thinner inner hyponeural band, both separated by a thin layer of connective tissue. The ectoneural system is furher organized in three layers: 1) an opaque and thin extracellular hyaline layer, covering the surface of the cells; 2) the somatic zone containing the cell bodies of the epithelial supporting (glial) cells plus the neurons; and 3) the neuropile, a fibrillar zone mainly composed by neurofibrillae intermixed with the apical parts of the supporting cells and containing intermediate filament bundles. In contrast, the hyponeural system is very thin, hardly distinguishable from the covering coelomic epithelium and in directly contact with the fluid of the hyponeural sinus. Serial sagittal sections of all samples at 1 day PA showed an incomplete RNC with a gap, corresponding to the ablated zone (**Fig 2c**). The ends of the injured nerve cord displayed an area of disorganized neuropil structure though they were already healed by a layer of cell bodies, secreting a thin hyaline layer (**Fig 2e**). On day 7 PA, and in the majority of the samples the nerve was no longer ruptured (**Fig 2f**), indeed, the gap was now filled with a very thin nerve (**Fig 2g**) lining the radial water channel. This nerve was characterized by the presence of layers of the cellular bodies located in the ectoneural and the hyponeural systems, although in the inner part of the filled gap the distinction between the two neural components was not possible, here the neuropil was also hardly visible. On day 14 PA, the neuropil was thicker and more organized than on day 7 and the tissue pattern of the regenerating RNC resembled that of uninjured RNC, although often with a reduced thickness (**Fig 2h** and **i**). Besides the RNC, other tissues/structures were locally affected during the post-ablation period. The podia at the level of the nerve gap remained retracted from day 1 to day 14 (**Fig 2a, c, f** and **h**), this feature being also visible by external inspection (see above). Additionally, in all regeneration samples a localized area of hypertrophy of the hyponeural sinus and of the radial water channel (**Fig 2d**) was evident. The swelling of this coelom-derived canal, with the associated hemal sinus, was particularly evident at day 14 PA.

**Fig 2.**
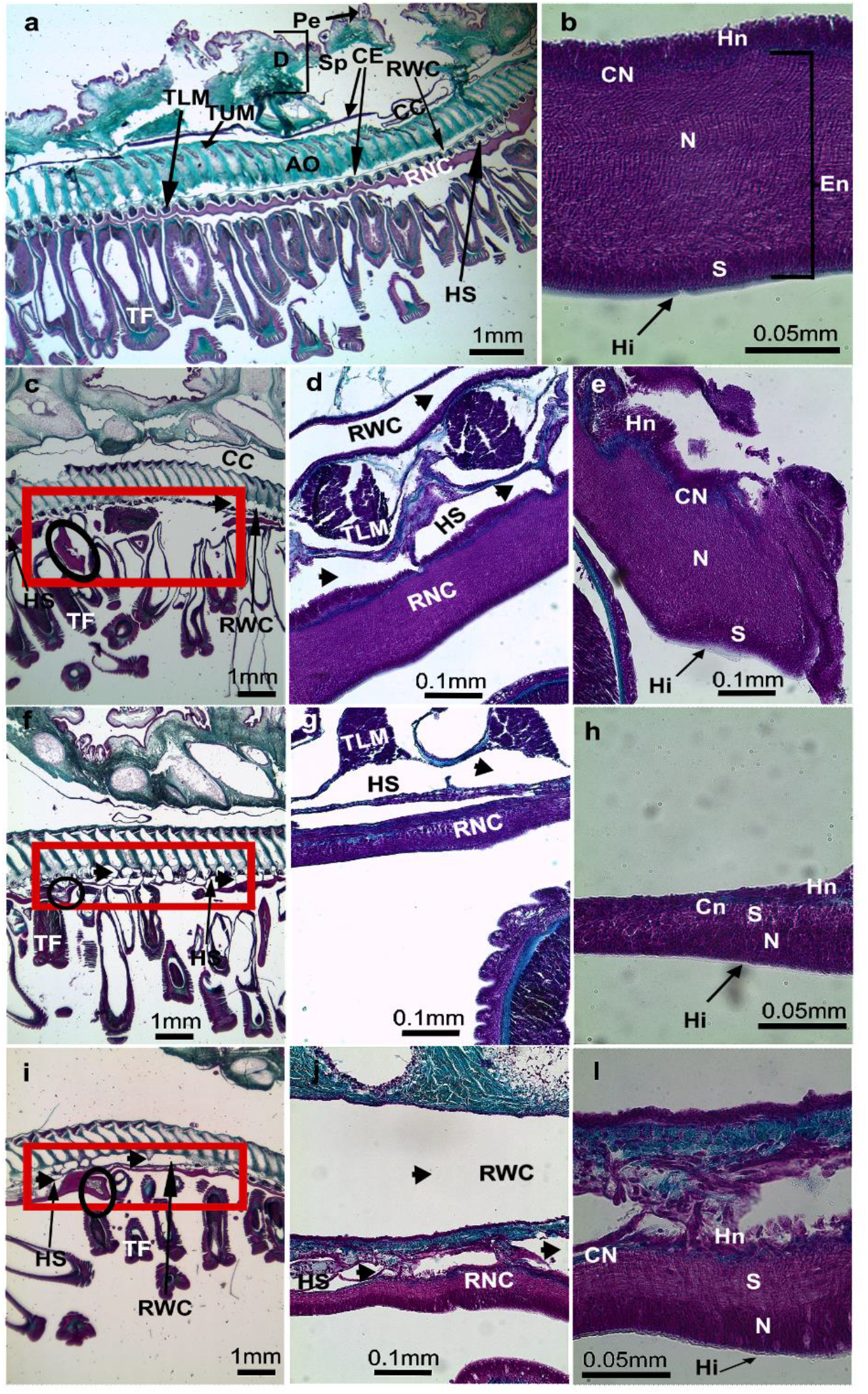
Sagittal sections of the starfish arm in control conditions (**a** and **b**), in day 1 PA (**c**) with the hypertrophy of the coelomic canals (arrow heads) (**d**) and the lump at the end of the injured RNC (**e**), in day 7 PA (**f** and **g**), and in day 14 PA (**h** and **i**). Orientation: the proximal part in the left side to distal part on the right side; aboral side at upper limit and oral side at the lower limit. **AO**- ambulacral ossicle; **Cn**- connective tissue; **CC**- coelomic cavity; **CE**- coelomic epithelium; **D**- dermis; **En**- ectoneural; **Ep**- epidermis; **Hi**- hyaline layer; **Hn**- hyponeural; **HS**- hyponeural sinus; **N**- neuropil; **Pe**- pedicellariae; **RNC**- radial nerve cord; **S**- somatic zone (cell body layer); **Sp**- spine; **TF**- tube foot; **TLM**- transversal lower muscle; **TUM**- transversal upper muscle. Representation of the radial nerve cord: (circle) the lump at the non-regenerative end, and (square) the nerve gap/tissue in regeneration.

### Dynamics of coelomocytes populations

Coelomic fluid circulating coelomocytes populations were characterized during RNC regeneration by FC. The characterization of the cellular populations was made based on the DRAQ5 intensity and Side Scatter (SSC)-A values. This allowed us to detect two distinct populations (P1 and P2) with different morphological complexity and fluorescence properties, in non-regeneration conditions (**Fig 3a**), as reported in our recent publication (Andrade et al. 2021). Under nerve regenerating conditions, a similar profile was detected on day 1 PA (**Fig 3b**). Interestingly at day 7 PA, a new coelomocyte population, here designated as P3, was detected in some animals (**Fig 3c**). At day 14 PA, P3 was already present in all animals (N=5) in a higher amount (**Fig 3d**). This new cellular population tends to display intermediate values of DRAQ5 incorporation in P1 and P2 cells, however apparently with side scatter values similar to those of P1 cells, suggesting similar morphological characteristics to this, latter, cell type (**Fig 3c** and **d**). It is important to notice that cells detected outside the limits of the three populations displayed in Figure 3 were not placed in a sufficient abundance zone of the plot and thus they are not considered another population; identifiable also as cell aggregates.

**Fig 3.**
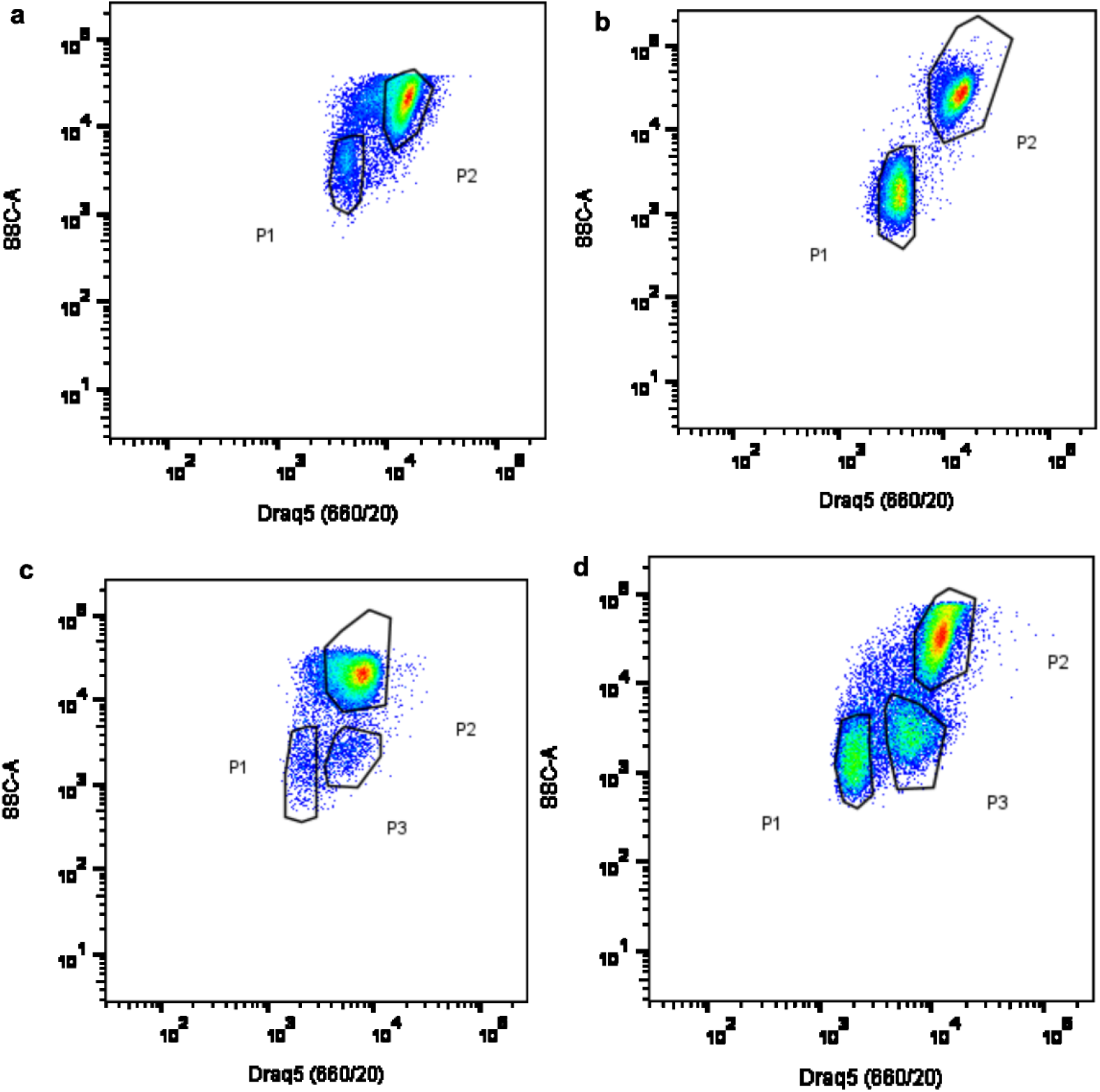
Coelomocytes flow cytometry analysis during Radial Nerve Cord (RNC) regenerative process. Circulating coelomocytes were examined by looking at Median Fluorescence Intensity of DRAQ5 vs SSC-A on the control (**a**), and days 1 (**b**), 7 (**c**) and 14 (**d**) after partial nerve excision. N= 6 controls and 5 per each time-point.

The percentage of cells of each population detected in the regenerative and non-regenerative samples was also determined. These values for the P1 and P2 populations do not change during the nerve regeneration process (**Fig 4a** and **b**). At days 7 and 14 PA the number of cells of the new detected population (P3) (**Fig 3c** and **d**) showed a significant increase (p<0.05) as compared to the values measured in day 0 or 1 (p<0.05) (**Fig.4c**).

**Fig 4.**
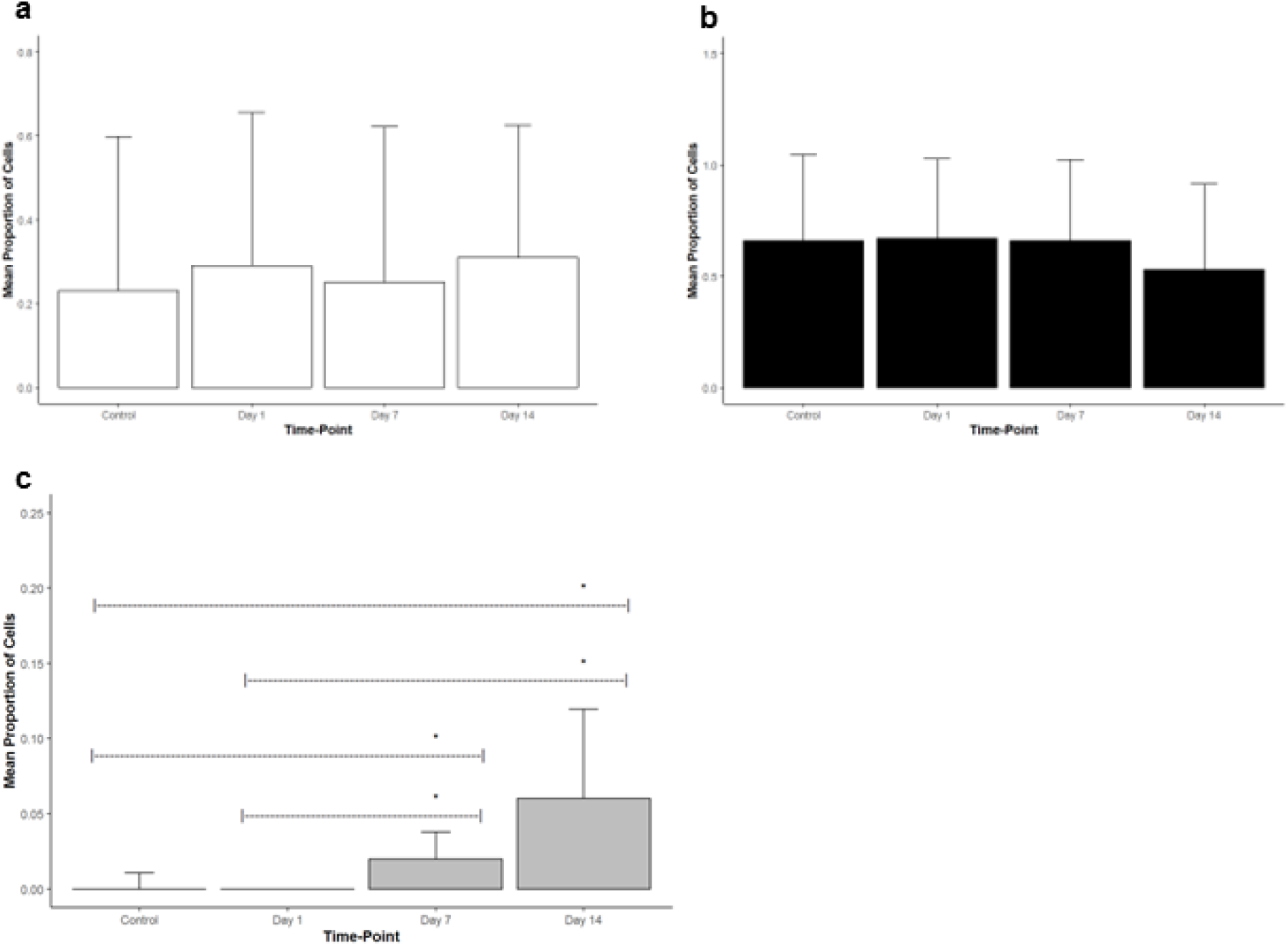
Bar plots of the mean percentage of P1 (**a**), P2 (**b**) and P3 (**c**) cells in control and at each time-point (1, 7 and 14 days) after partial radial nerve ablation. (N=6 controls and N=5 specimens per each time-point). * Significant differences were determined (p<0,05).

### Characterization of coelomocytes populations and of their morphotypes through IFC

In accordance with the FC results, IFC of circulating coelomocytes was performed during RNC regeneration, in order to further characterize morphologically the new P3 cells. Before the analysis, the focused single cells were selected in the IDEAS software using the parameter “root mean square gradient” (RMS=40-80) and the relation between “aspect ratio” and the area of the cells in the bright field (BF) channel. Applying similar FC selective parameters, intensity in SSC channel and intensity of DRAQ5, it was possible to discriminate P3. The cells of this population show a circular/oval shape with some cytoplasmic extensions, displaying an outer bright (light or dark) periphery, containing inside several dark granules (**Fig 5a**). Matching the FC results, P3 cells display DRAQ5 intensity values between those of P1 and P2 cells, a similar SSC values to the P1 cells (**Fig 5b** and **c**). According to our FC data, a low number of P3 events (8-11%) was detected in the coelomic fluid of non-regenerating starfish (**Fig 5b**). At 14 PA (**Fig 5c**), though a significant increase in the number of P3 cells was determined (22-30%).

**Fig 5.**
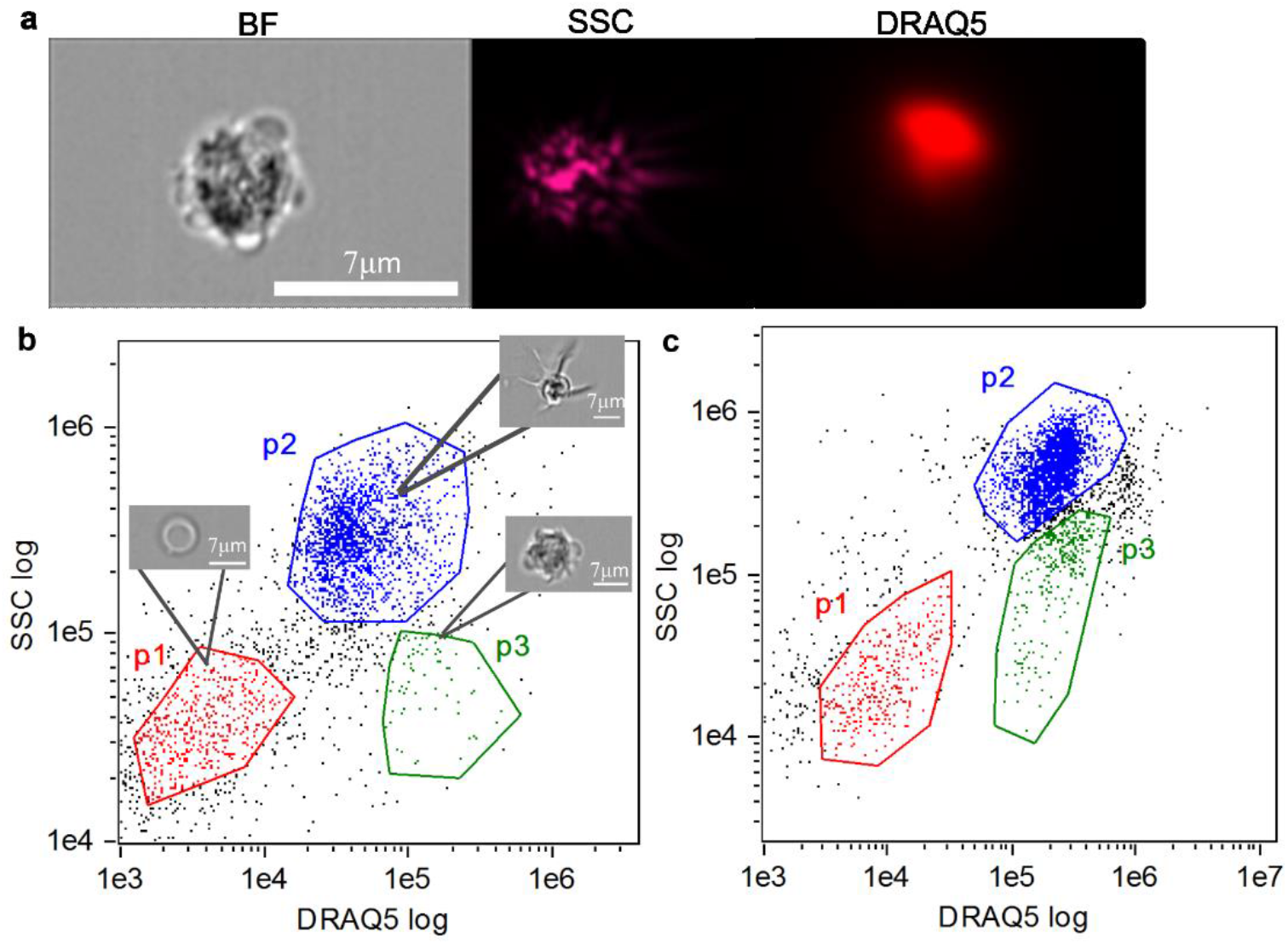
IFC characterization of the P3 population stained with DRAQ5. (**a**) Bright field image and DRAQ5 fluorescence image. (**b**) Dot plot showing cell area vs DRAQ5 intensity under non-regenerating conditions. (**c**) Dot plot showing cell area vs DRAQ5 intensity at 14 days post-partial nerve ablation. Red, green and blue dots correspond to P1, P2 and P3, respectively.

A selection of the circulating coelomocytes populations and of their morphotypes was also performed using the IFC parameters, in order to obtain an efficient method to choose each cell type. The better discriminatory features were defined using an artificial intelligence (IA) algorithm, namely the wizard “Feature Finder” of the “Guided Analysis” within the IDEAS software starting with the focused single cells attributed to each population as training set. **Table 3** presents the discriminatory parameters for the several populations and their respective morphotypes. According to this algorithm, P1 is discriminated from P2 by the cell area and height. The P3 cells differ from P1 cells in terms of DRAQ5 signal intensity and BF size features and differ from P2 cells mostly in BF size features. The two P1 morphotypes previously described by fluorescence microscopy (Andrade et al. 2021) were also discriminated by the IA algorithm. Here, the P1S morphotype presents a smaller nucleus-to-cytoplasm ratio and a heavily granulated cytoplasm, surrounded by an uninterrupted bright red membrane. Nuclei of the P1L morphotype occupy most of the cell space that show a less heterogeneous BF image. In addition, and by IFC, a third P1 morphotype was detected and was designated as P1 Brighter cell (P1B). This morphotype can be described as having a high nucleus to cytoplasm ratio and is contoured by a bright halo (**Fig 6**). Regarding the P2 population, the phagocytes (petaloid vs filopodial morphotypes) differ in terms of diverse BF features related to cell shape. The thrombocytes (regular vs big granulated morphotypes) can also be differentiated through some BF features related to cellular texture (i.e.: cellular internal complexity). Both phagocytes type (petaloid vs filopodial) can be further differentiated from both thrombocytes morphotypes (polygonal vs big granulated) through BF features of cell shape. These differentiations were also done by fluorescence microscopy as described in Andrade et al. 2021..

**Table 3.**
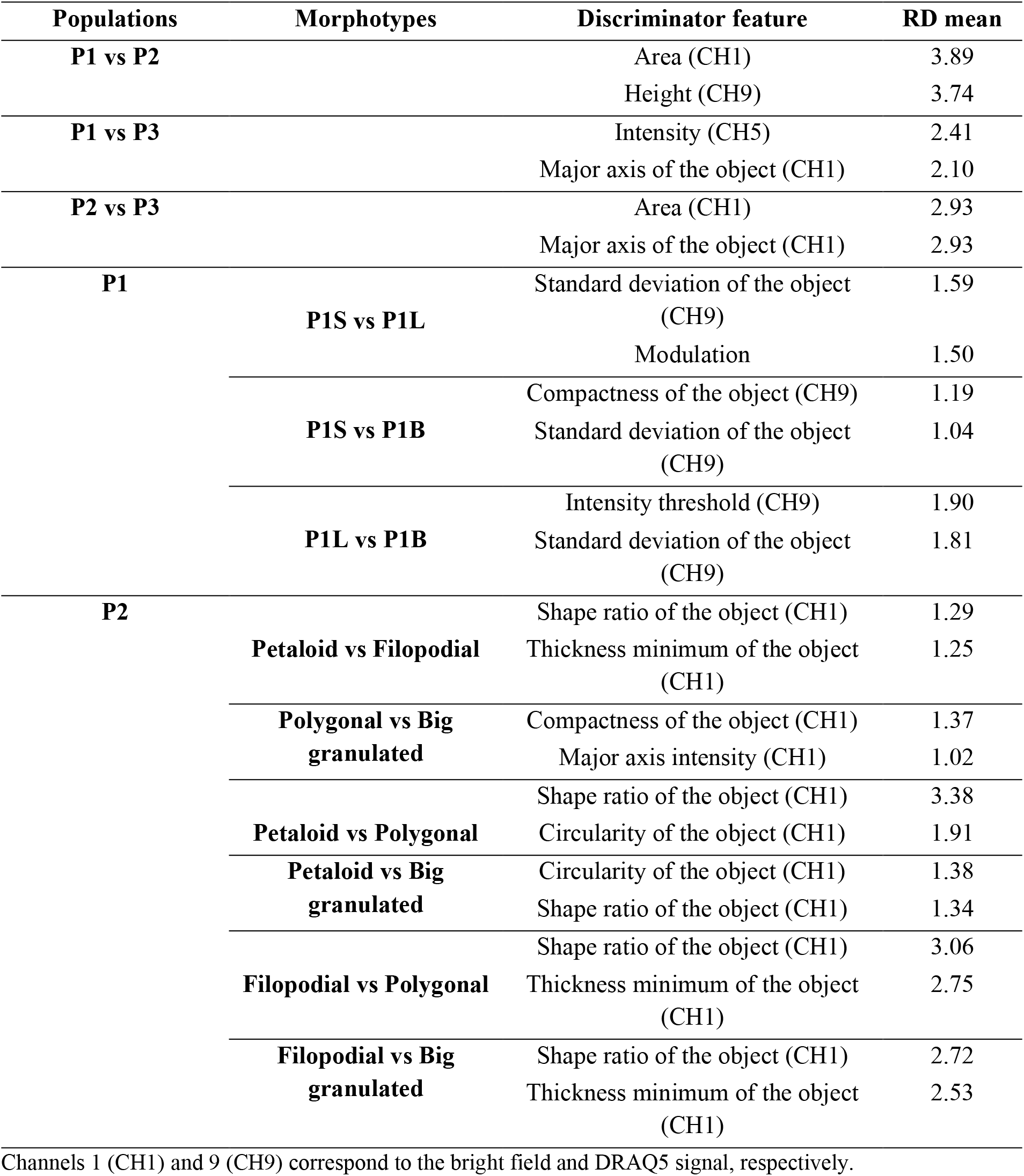
Discriminatory features for P1, P2 and P3 populations, and of their respective morphotypes, P1S, P1L, P1B, and petaloid, filopodial, polygonal and big granulated.

**Fig 6.**
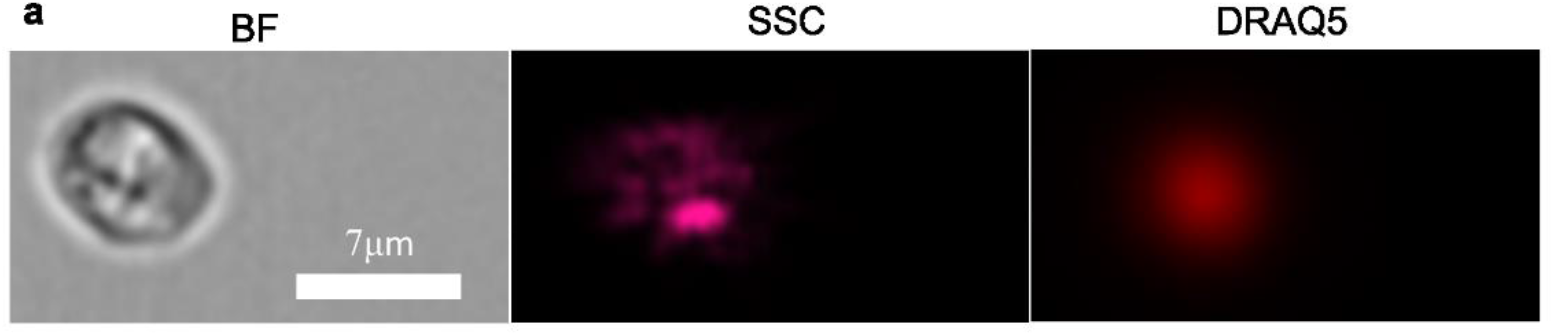
Bright field image and DRAQ5 signal intensity of the new P1B morphotype.

The three P1 morphotypes can be localized in the IFC dot plot through a 2-step method after applying the discriminatory IFC features to P1 cells (**Fig.7**). First, are selected the P1L and P1S cells morphotypes through their best discriminatory features (**Fig.7a**). Then, the P1B cells, discriminated from the other P1 morphotypes through the parameters “modulation” in (CH1) and “standard deviation of the object” in (CH9), are added (**Fig.7b**).

**Fig 7.**
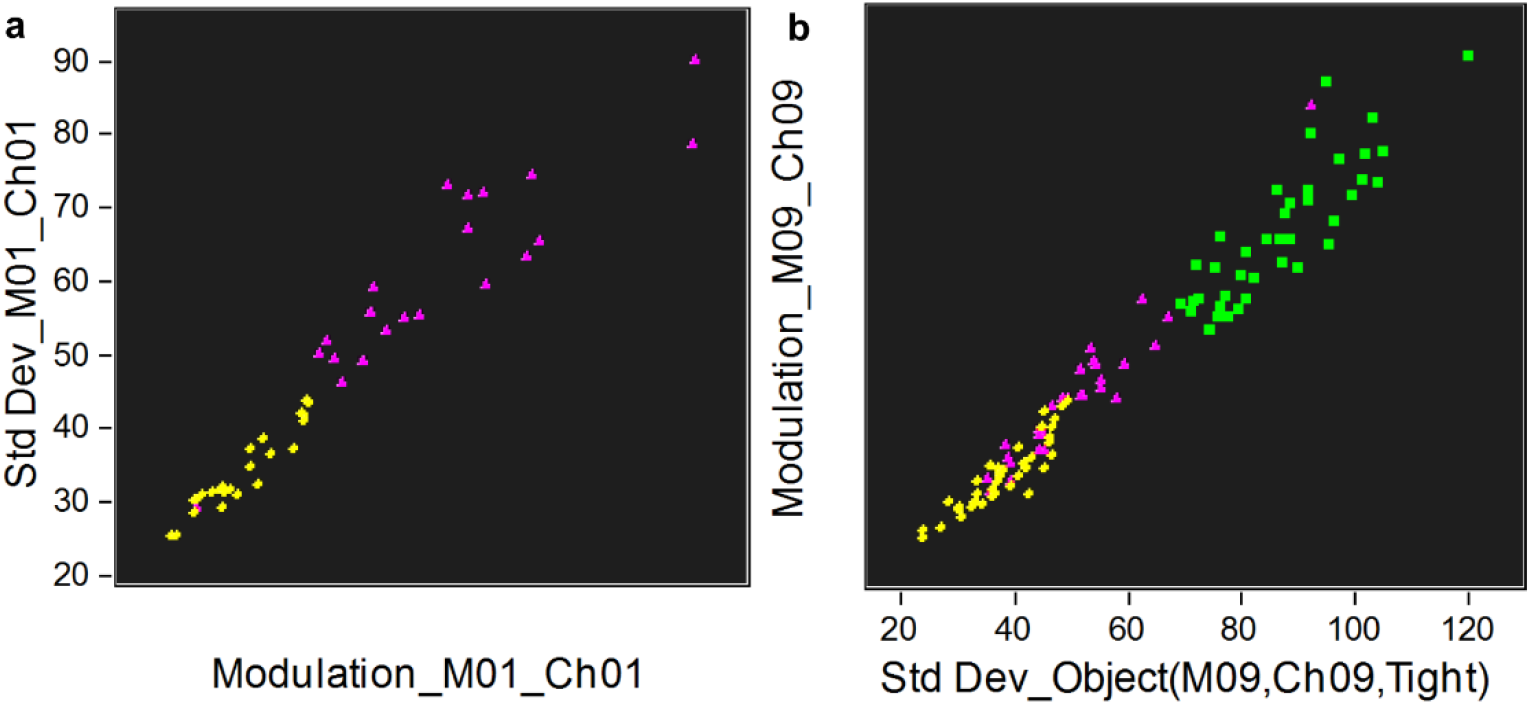
IFC discrimination of the morphotypes from P1. (**a**) Dot plot of the differentiation between P1L and P1S through the relation between texture features, and (**b**) between P1B with P1S and P1L cells. Yellow diamond- P1L; green square- P1B; purple triangle- P1S.

Regarding the P2 morphotypes, a 3-step method can be used with the same purpose. The thrombocytes and phagocytes can be discriminated through the selection of the parameters: “shape ratio” in (CH1) and “circularity of the object” in (CH1). In particular, phagocytes and thrombocytes are selected through a gate range of 0-0.6 and 0.8-1.0 for “shape ratio” and of 0-10 and 20-50 for “circularity of the object”, respectively (**Fig 8a**). Further, the two morphotypes of each phagocyte and thrombocyte can be selected trough their best discriminatory features, as specified in Table 3 (**Fig 8b** and **c**).

**Fig 8.**
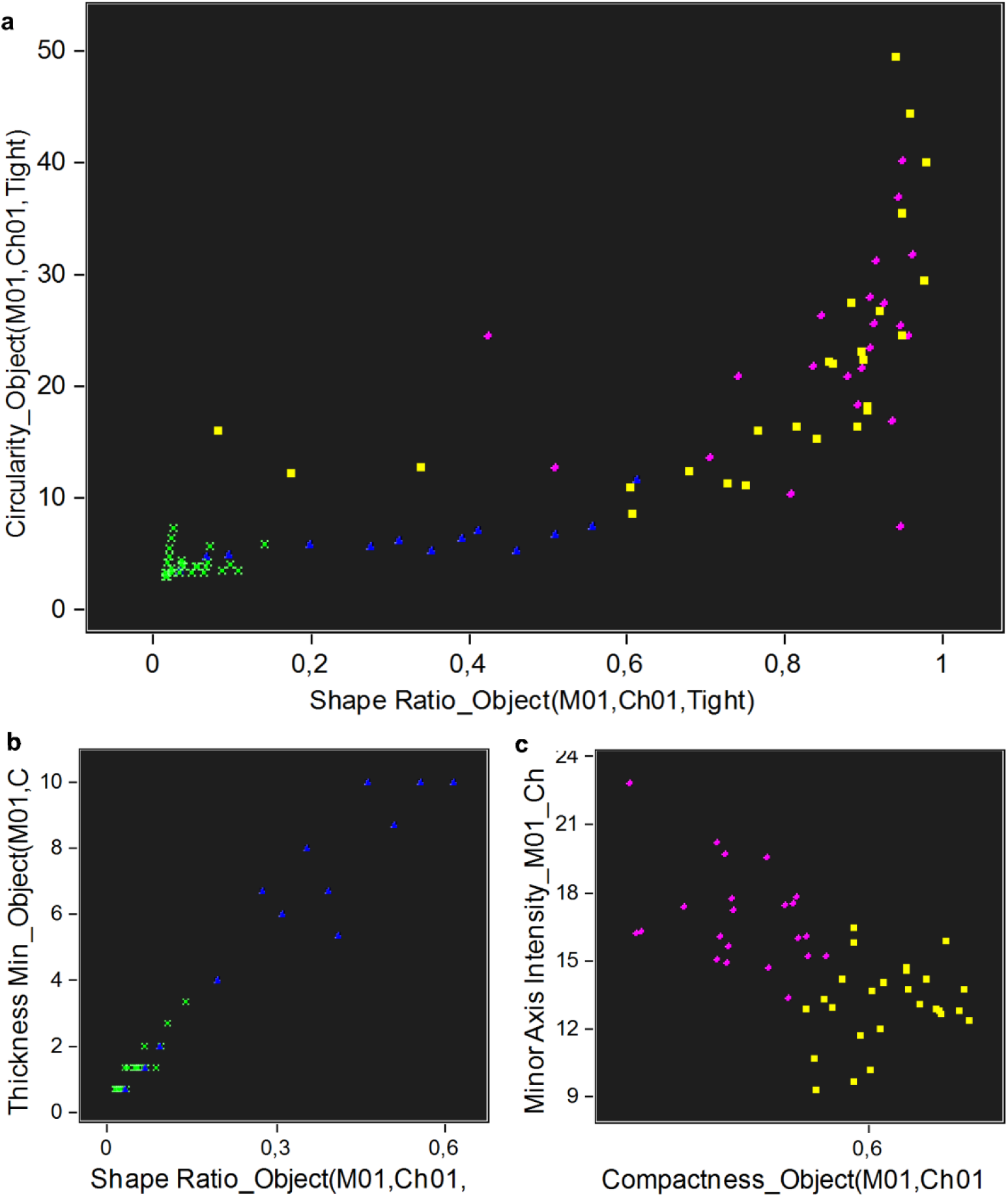
IFC discrimination of the morphotypes from P2 (**a**) Dot plot of the differentiation between phagocytes (petaloid and filopodial cells) and thrombocytes (polygonal and big granulated cells, (**b**) petaloid filopodial cells, and (**c**) polygonal and big granulated cells, through the relation between cell shape features. Green cross-petaloid; blue triangle-filopodial; yellow square-big granulated; purple circle-polygonal.

## Discussion

### Nerve ablation triggers changes in the starfish mobility behaviour and creates anatomic anomalies in the regenerating arm

Animal behavior comprises two different but complementary mechanisms: innate and adaptive. The adaptive behavior was defined by Migita and colleagues (Migita et al. 2005) as the capacity of the animal to flexibly change its innate behavior pattern when it is put into a situation where this does not serve its purpose. Since starfish lack a self-awareness or consciousness complex system, a self-organized system, such as neuromuscular activities, could be the answer for the modulation of their innate behavior (Kelso 1995). Starfish CNS is connected to various parts of the starfish body by peripheral motor nerves of the peripheral nervous system (PNS). The RNC extensively innervates several effectors of the starfish arm, like the tube feet, muscles, peritoneum, epidermis, and numerous elements of the mutable collagenous tissue, through metameric nerves (Mashanov et al. 2016). Even though it has not been functionally demonstrated that the RNC controls these effectors, former literature has reported that certain tracts of the RNC have motor functions and might be specifically dedicated to the orientation of the animal (starfish) during locomotion. Moreover, the RNC might transmit local information to the five arms in order to produce a coordinated behavior through the nerve extensions (Dale 1999; Kerkut 1954). Although, a self-organization of the tube feet alone has, probably, the potential to change the starfish movement (Migita et al. 2005). Starfish choose one arm to lead their movement and when they are presented with an obstacle, they can modify their behavior by changing the leading arm chosen or simply by helping with the original arm with the recruitment of the adjacent arms. Therefore, this behavior could be due to a multi-layer mechanism of self-organization including a lower self-organization of the tube feet (Migita et al. 2005).

In this present study, the partial ablation of the RNC has triggered changes in the starfish displacement (arm leading of the animal movement and loss of podia activity) and anatomic modifications in the injured arm (podia retraction, local bending of the arm and misalignment of the lateral spicules). These changes remain for most of the regeneration period. Assuming that the CNS commands the peripheral innervated effectors, the partial excision of the RNC tissue could have caused their loss of functionality, and consequently may have directly affected the podia and spicules movements and, thus, indirectly the local arm bending. On the other side, after starfish arm tip amputation the first emergency reaction is characterized by a strong and rapid body wall contraction at the injured site (Ben Khadra et al. 2015a; Ben Khadra et al. 2017). This process was observed to be associated with the partial contraction of the muscle layers of the perivisceral coelom (the main body cavity), probably acting as a defense reaction to reduce the loss of body fluids (Ben Khadra et al. 2017). Moreover, it was observed that the contraction pulled the podia towards the central area of the wound (Ben Khadra et al. 2015a). Therefore, podia retraction observed by the analysis of external anatomy and of the histological sections (**Fig 1** and **2**), could be associated to a similar muscle contraction process, or to the loss of innervation. The retraction of the podia that caused their inactivity between days 1-14 PA, hence, could also be a defense mechanism to protect the wounded nerve. Starfish suffered a change in the use of the NR as leading arm of the movement after the partial excision of the RNC, failing to use the NR and overly using the NRA. Indeed, starfish have a deterioration in appetitive response towards the prey when the leading arm is the one with the nerve in regeneration (Piscopo et al. 2005). The authors of this paper claimed that the cause behind this behavior was the loss of the neuronal system tissue dedicated to the motor function, since starfish recovers the appetitive response when this neuronal system is regenerated (Piscopo et al. 2005). Nonetheless, the loss of the RN use as leading arm between days 1-9 PA could be a be a byproduct of the retraction and inactivity of the podia. Interestingly, between days 9 PA and 14 PA the RN had a weak increase in its use as leading arm, which coincides with the regeneration period. A fully recovery is seen when the nerve is fully regenerated and displaying a similar morphology and structure to the non-regenerating nerve tissue (the reference). Moreover, when the starfish displaces and changes its direction, since the RN arm podia are inactive only at the site of lesion, the zone of the arm away from the injury still performs movement in the direction of the leading arm, which the zone with the injury does not follow. These contrasting effects in terminal and proximal parts of the injured arm cause a bending of the RN arm tip (**Fig 1**). Although, as seen in Ben Khadra et al. 2015a, a complementary mechanism may be operating, since the strong contraction of the starfish body wall could be also causally related to the generation of this anatomical anomaly. The latter is due to the fact that during the arm’s tip regeneration the body wall shows a particularly strong constriction only at the distal portion of the injury stump. An additional effect is observed over the spines and spicules, which in starfish are known to be mechanoreceptive (Garm 2017). Their observed misalignment (**Table 3** and **Fig 1**) could be a possible consequence of deficient nerve tissue coordination. In summary, the main causes for the adjustments of animal anatomy and of its displacement because of the partial loss of CNS tissue, may be the lack of connection of its motor behavior, due to the partial loss of CNS tissue, may be the result of a lack of functional connections between the innervated effectors and the body wall contraction.

The appearance of the oedema in the wound area (at 1-2 PA for 50 and 65% assayed animals) can be seen as a first emergency reaction, a swelling of the tissue that is usually associated with injury and results in a fluid buildup. This phenomenon was observed previously during starfish arm tip regeneration, between 72 h and 1 weak post-amputation (Ben Khadra et al. 2017). During arm tip regeneration, the oedema is referred to a temporary area of granulation tissue composed by fibroblasts, phagocytes, nervous elements, dedifferentiating myocytes and undifferentiated cells embedded in a highly disorganized collagen/ECM matrix, which is placed just below the epidermis and serves as wound protection (Ben Khadra et al. 2017). However, signs of this process were not observed in our histological examinations, eventually since it could be mainly associated only with the enlargement of the coelomic cavity due to fluid accumulation.

### Histology reveals insights of the arm tissue patterns during RNC regeneration

Histological analyses of arm samples allowed to define the tissue pattern of recovery of the injured RNC as well as those of the surrounding tissues during nerve regeneration. All echinoderms, like the starfish possess a so-called CNS lacking a centralized brain. This neural system is characterized by the presence of a circumoral nerve ring in the central disk and five radial nerve cords (RNC) that run along the arms, in parallel to the radial water canal (Ruppert et al. 2004; Mashanov et al. 2016). The RNC consists of an agglomeration of neurons and radial glial cells associated with an extensive neuropil (Mashanov et al. 2013). The radial glial cells are non-neuronal cells of the nervous system and are responsible for the support and homeostasis of the neurons (Ortega and Olivares-Bañuelos 2020). As previously noted, the herein described RNC has a similar composition, morphology, and structure to that of other reported asteroid species (Hymann 1955; Ben Khadra et al. 2015b), in which the RNC has a V-shaped morphology, it is continuous in its lateral sides with the epidermis, and whose main component is the ectoneural system. The same tissue organization was reported, for instance, in the Holothuroidea (Mashanov et al. 2013).

Histologic studies on starfish nerve regeneration have been mainly focused on the arm tip regeneration that followed by a traumatic lesion (Moss et al. 1998; Mladenov et al. 1989; Fan et al. 2011; Ben Khadra et al. 2015a; Ben Khadra et al. 2015b; Ben Khadra et al. 2017). This process usually involves three main phases: 1) a repair phase, characterized by the wound healing, which involves an accumulation of phagocytes and thrombocytes-like cells at the injury site to protect the inner body environment, and by the first signs of neurogenesis; 2) an early regenerative phase, during which tissue reorganization and first signs of tissue regeneration phenomena occur; 3) an advanced regenerative phase, characterized by tissue restoration and regrowth with the formation of a new small regenerating arm consisting of the same structures of the adult arm (Ben Khadra et al. 2015a; Ben Khadra et al. 2015b; Moss et al. 1998). This multistep process can be seen also in the regeneration of the RNC after partial ablation (Ben Khadra et al. 2015a). This process is followed by our samples. In fact, when compared to observations made in the starfish *Echinaster sepositus* (Ben Khadra et al. 2015a), the nerve end is healed at 1 day PA. This healing is characterized by the re-epithelization of the RNC wound margins, which is carried out by cells, at least partially deriving from the nearby nerve stumps, although a contribution from more distant migrating/circulating elements cannot be excluded. Therefore, in echinoderms the timing of the repair phase is apparently maintained, regardless of the species analysed. Moreover, the structure/tissues involved in the stump healing seem to be identical (Ben Khadra et al. 2018). In the second phase, at day 7 PA, the neuropil zone starts to differentiate. This pattern again perfectly resembles that of *E. sepositus*; in the latter the supporting radial glial cells are the first component to be formed in the new neural tissue, where they rebuild the structural network needed for the subsequent repopulation of neurons (Ben Khadra et al. 2015b). Further similarities can be found in the regeneration of the RNC after partial nerve ablation of the sea cucumber *Holothuria glaberrima* (Miguel-Ruiz et al. 2009). An interesting aspect is that cell proliferation is not the main mechanism in holothurian RNC regeneration. Moreover, there is evidence that RNC regeneration does not rely upon resident undifferentiated/stem cells (Mashanov et al. 2014). Instead, the mechanism behind the formation of new neural cells probably occurs through a mechanism that includes dedifferentiation of the nearby radial glial cells and the recruitment of other cells migrating to the area possibly from deeper regions of the neuroepithelium (Mashanov et al. 2017; Mashanov and Zueva 2019). Therefore, radial glial cells serve as a source of new neuronal elements and new glial cells all contributing to the regrowth of the nerve tissue (Mashanov and Zueva 2019). The last phase, at day 14 PA, corresponds to the growth and terminal differentiation of the newly regenerated tissue, until it reaches the same size and structural organization of the non-regenerating tissue.

### A new coelomocyte population was detected trough FC during RNC regeneration

Besides the recovery of the neural tissue itself, as demonstrated for Holothuroidea (Mashanov et al. 2017), the involvement of other tissues surrounding the injured RNC is a further important aspect to consider. Indeed, both the traumatic event of nerve ablation and the subsequent tissue regrowth and remodeling are expected to cause local, but possibly also systemic, inflammatory/immune responses. This is particularly true for all the coelomic and coelomic-derived cavities, in which the typical immune cells of echinoderms, the coelomocytes, normally circulate. As expected, a localized hypertrophy of the hyponeural sinus (which directly faces the RNC) and of the radial water canal was present during the whole period of regeneration. Similarly, a moderate hypertrophy was observed in the nearby haemal sinus, a non-coelomic system of lacunae also involved in immune functions, that contain coelomocytes as well. This phenomenon could be explained by the overuse of the coelomic/coelomic-derived canals for the transport of both molecules and immune or other migrating cells from the neighbor tissues, as suggested by other authors (Hernroth et al. 2010; Guatelli et al. 2022). The changes in coelomocyte composition that we found during the period of RNC regeneration and discussed below are perfectly in line with this idea.

The cytotypes, and most importantly, the origin of cells that circulate in the echinoderm coelomic fluid have been a long matter of discussion. Recently, our research (Andrade et al. 2021), resorting on the use of flow cytometry (FC) plus an imaging combined approach, allowed us to characterize minutely two populations of coelomocytes in starfish, denominated as P1 and P2. These two cell populations differ substantially in abundance, cell size, cell morphological complexity, and the incorporation of fluorescent DNA intercalator. The P2 cells displayed higher side scatter (SSC) and forward scatter (FSC) values than P1 cells, probably due to a more complex/structured surface and a richer internal cytoarchitecture. Moreover, P2 cells incorporate approximately double of the amounts of DRAQ5 and are more abundant than P1 cells, representing 60-70% of the total circulating coelomocytes.

These two coelomocyte populations presenting the same FC features were also identified in the present study. These populations are present in similar proportions in all experimental groups, both in the control condition and in the nerve regenerating samples (at all time-points). When analyzed previously (Andrade et al. 2021), two morphotypes of the haploid and low mitotic active P1 cell population were detected. One with a less heterogeneous nucleus and a higher nucleus-to-cytoplasm ratio, was hypothesized to be a non-differentiated form of the more mature morphotype. This latter had a lower nucleus-to-cytoplasm ratio and contained a heavily granulated cytoplasm. The other, P2, population was described in the same publication. Two main morphotypes were revealed in it: one similar to phagocytes, designated as petaloid and filopodial cells, and another referred as thrombocytes-like and constituted by regular (polygonal) and big granulated cells. Additionally, two other P2 morphotypes were observed only by fluorescence microscopy. In the present study a new population, designated here as P3, was detected among the circulating coelomocytes between days 7 and 14 PA, and was futher characterized through the use of FC and IFC methodologies. P3 cells have similar internal complexity than P1 cells and show a DRAQ5 incorporation with intermediate values between those of P1 and P2 cells.

The origin of the P3 coelomocyte remains unknown. The axial organ (Leclerc and Bajelan 1992), the Tiedemann’s bodies (Kaneshiro and Karp 1980), the hemal organs (Ferguson 1966), and the coelomic epithelium (Guatelli et al. 2022; Sharlaimova et al. 2021; Bosshe and Jangoux 1976) were suggested as candidates for the so-called hematopoietic tissue in starfish. Nonetheless, this last tissue is, nowadays, the most accepted tissue candidate to carry out the mentioned functions. A particular relevant issue here is understanding the source of circulating coelomocytes, and their constituent populations. Initial microscopy analysis showed that the coelomic epithelial cells share similar ultrastructural features with the circulating coelomocytes (Holm et al. 2008; Gorshkov et al. 2009; Sharlaimova et al. 2010), in particular, with thrombocytes-like cells (Guatelli et al. 2022). Moreover, proteomic analysis showed the presence of some common proteins in the coelomic epithelium and the circulating coelomocytes of *M. glacialis*, including proteins related with the cell motility, which would suggest the migratory behaviour of cells from the coelomic epithelium, moving from this to the coelomic cavity. Most relevant to the regeneration process is the fact that histology has showed that coelomic epithelium cells undergo partial dedifferentiation and subsequent epithelial-mesenchymal transition during arm tip regeneration. This would point to the direct involvement of coelomic epithelial-derived cells to the rebuilding of regenerating structures. However, a direct release of free wandering coelomocytes from the apical part of the coelomic epithelium towards the coelomic lumen was never observed, either in non–regenerating or in regenerating conditions (Guatelli et al. 2022). Alternatively, and as pointed out above, coelomic epithelium could be involved in the hypertrophy of the coelomic and coelomic-derived canals by providing immune response through immune factors. In this context, the physiological role of the P3 population, though apparently relevant in the regeneration process, needs to be determined.

## Conclusions

This present study brings forward new insights into CNS regeneration in the starfish *M. glacialis*, focusing on the RNC post-traumatic regeneration. Approaching several methodologies in a complementary perspective, such as a behavioral assay and anatomical observations, histological analysis, and a combination of FC and IFC, it was possible to investigate starfish displacement, external body modifications and internal tissue patterns to sketch some of the cellular processes involved in RNC regeneration.

The partial excision of the nerve tissue of RN arm implicated an impact on the overall animal movement and arm coordination. However, when the missing nerve tissue was under the restoring process, the starfish began to recover the motor function of the disabled arm. The nerve lesion also led to external anatomic anomalies as a defense mechanism of the wounded area. The histology study provided a vivid view of the cellular processes and tissue patterns associated with nerve tissue regeneration. The most striking aspect was the resemblance of the tissue patterns during RNC regeneration with that described for the starfish arm regeneration (Ben Khadra et al. 2015a; Ben Khadra et al. 2015b). Interestingly, the somatic zone regenerated before the neuropil, which is strong evidence for the hypothesis that the radial glial cells are involved in the formation of new neuronal and new support cells (Mashanov and Zueva 2019; Mashanov et al. 2013) and, consequently, of the RNC tissue. IFC analysis allowed the characterization of a new P1 morphotype in regenerating and non-regenerating individuals. Here is described a 2-step and 3-step method that allows identification through IFC and IDEAS of P1 and P2 morphotypes, respectively. A novel coelomocyte population, with a proposed designation of P3, associated with nerve regeneration was detected and characterized by FC coupled with IFC. In that manner, we point out to the necessity for implementing new omics methodologies that would help characterizing the function and involvement of the P3 cells, plus of the coelomic epithelium, in the RNC regeneration. Nonetheless, we firmly believe that this present study provides strong evidence that the regeneration of starfish CNS not only relies on the radial glial cells as progenitor/precursor cells for neurons but also on specific cell populations, but also suggests that most probably the coelomic epithelium plays a regulatory role as hematopoietic/immune tissue and/or as source of regeneration-competent cells.

## Supplementary information

Gating strategy of FC results firstly passed by the selection of all the positive events in the DRAQ5 channel that led to the exclusion of dead cells and high auto-fluorescent cells. Then, the relation between SSC-H (height) and FSC-A (area), and between SSC-A (area) and SSC-H was used to select the single cells and remove the aggregates (Supplementary figure 2).

**Supplementary figure 1.**
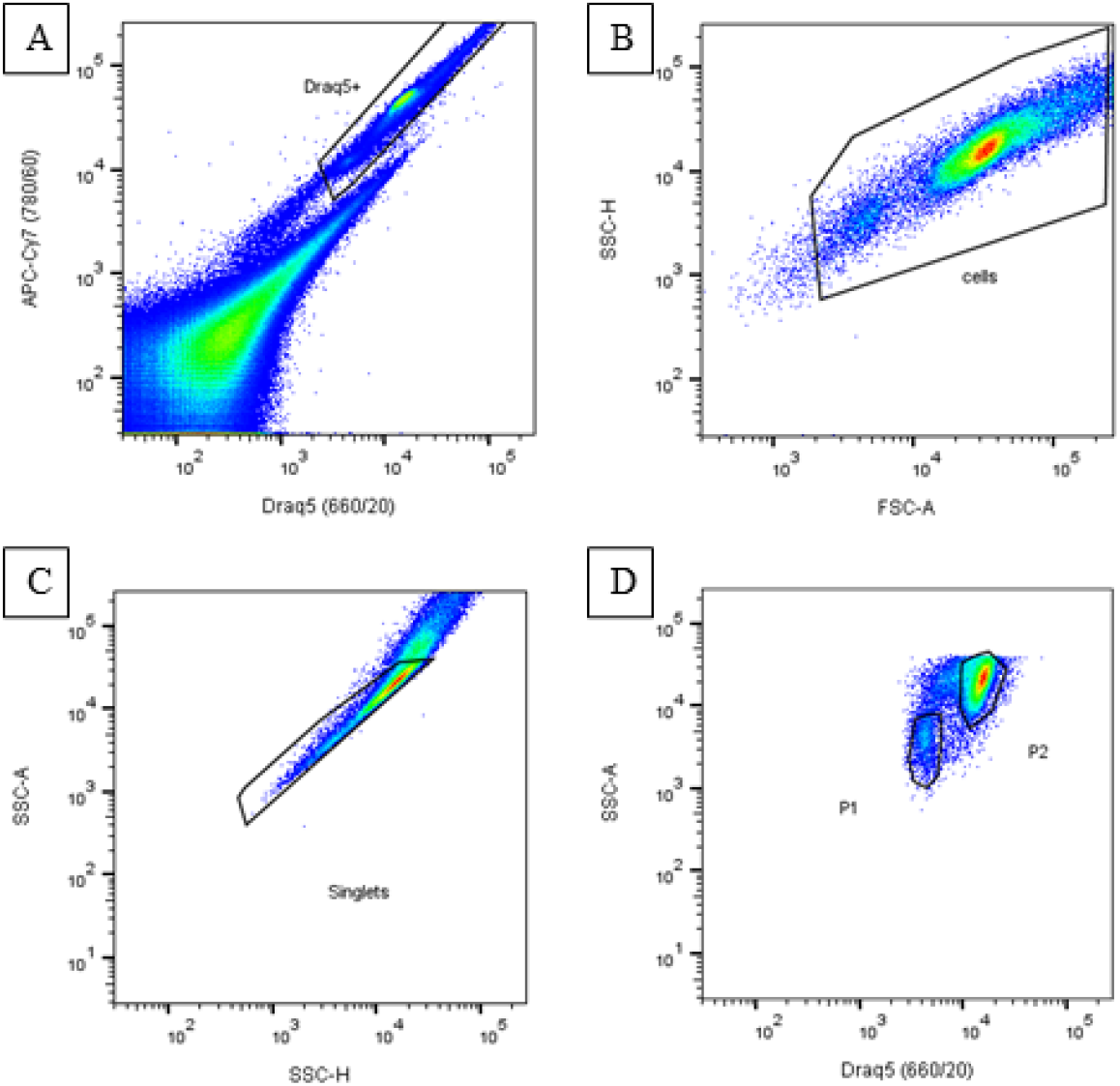
Representative staining and gating strategy of totally circulating coelomocytes. Cell population was selected by successive gating: (A) exclusion of high auto-fluorescent cells through the selection of DRAQ5 labeled cells; (B) exclusion of debris and some dead cells through the relation between SSC-H (height) and FSC-A (area); (C) removal of aggregates and selection of single cells through the relation between SSC-A and SSC-H and (D) P1,P2 and P3 populations selection by looking at median fluorescence intensity in DRAQ5 channel vs SSC-A.

