## Supplementary material for "Regeneration of Starfish Radial Nerve Cord restores animal mobility and unveils a new coelomocyte population"

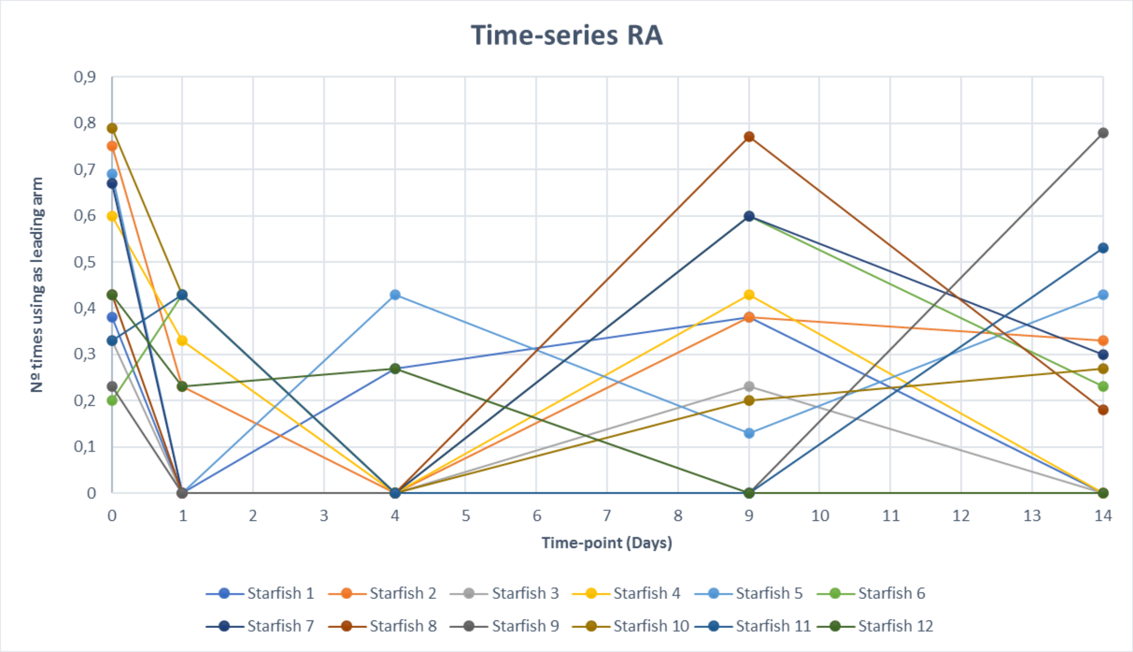


Supplementary Figure 1. Time series plot of the RN usage as leading arm by each starfish.

### Supplementary information

Gating strategy of FC results firstly passed by the selection of all the positive events in the DRAQ5 channel that led to the exclusion of dead cells and high auto-fluorescent cells. Then, the relation between SSC-H (height) and FSC-A (area), and between SSC-A (area) and SSC-H was used to select the single cells and remove the aggregates (Supplementary figure 2).

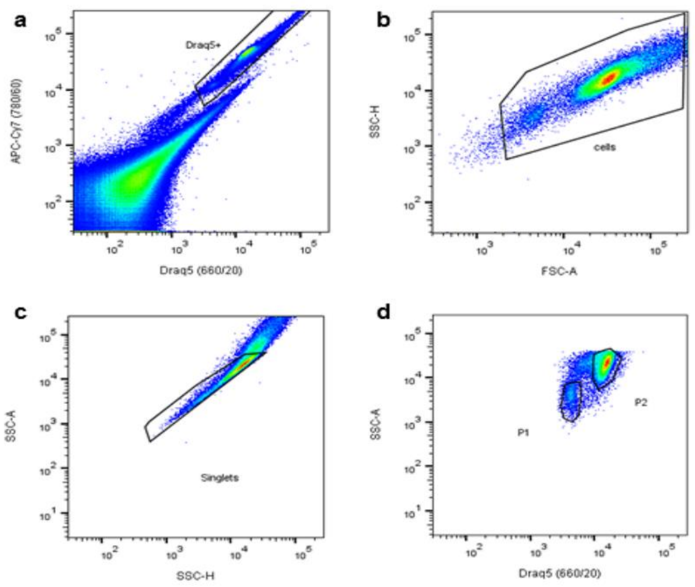


Supplementary figure 2. Representative staining and gating strategy of totally circulating coelomocytes. Cell population was selected by successive gating: (a) exclusion of high auto-fluorescent cells through the selection of DRAQ5 labeled cells; (b) exclusion of debris and some dead cells through the relation between SSC-H (height) and FSC-A (area); (c) removal of aggregates and selection of single cells through the relation between SSC-A and SSC-H and (d) P1,P2 and P3 populations selection by looking at median fluorescence intensity in DRAQ5 channel vs SSC-A.
